# Epigenomic plasticity of Arabidopsis *msh1* mutants under prolonged cold stress

**DOI:** 10.1101/263780

**Authors:** Sunil Kumar Kenchanmane Raju, Mon-Ray Shao, Yashitola Wamboldt, Sally Mackenzie

## Abstract

Dynamic transcriptional and epigenetic changes enable rapid adaptive benefit to environmental fluctuations. However, the underlying mechanisms and the extent to which this occurs are not well known. *MutS Homolog 1* (*MSH1*) mutants cause heritable developmental phenotypes accompanied by modulation of defense, phytohormone, stress-response and circadian rhythm genes, as well as heritable changes in DNA methylation patterns. Consistent with gene expression changes, *msh1* mutants display enhanced tolerance for abiotic stress including drought and salt stress, while showing increased susceptibility to freezing temperatures and bacterial pathogen *P syringae*. Our results suggest that chronic cold and low light stress (10 °C, 150 μE) influences non-CG methylation to a greater degree in *msh1* mutants compared to wild type Col-0. Furthermore, CHG changes are more closely pericentromeric, whereas CHH changes are generally more dispersed. This increased variation in non-CG methylation pattern does not significantly affect the *msh1*-derived enhanced growth behavior after mutants are crossed with isogenic wild type, reiterating the importance of CG methylation changes in *msh1*-derived enhanced vigor. These results indicate that *msh1*methylome is hyper-responsive to environmental stress in a manner distinct from the wild type response, but CG methylation changes are potentially responsible for growth vigor changes in the crossed progeny.

## INTRODUCTION

Plants have developed mechanisms to overcome constantly changing environments. Species that are more adaptable to changing environments through phenotypic plasticity and selection of adaptable traits survive. These changes occur at different levels, from morphological and physiological changes to modulations in gene expression and chromatin behavior, allowing plants to cope with the challenges of nature. While a major source of this adaptive response can be attributed to genetic variation (Franks and Hoffmann, 2012), recent studies are pointing towards the potential role of chromatin modifications and epigenetics in plant responses to environmental changes (Bilichak and Kovalchuk, 2016). Environment-induced epigenetic modifications are generally transient, and the consistency of the environmental cue perceived by plants plays a role in inducing epigenetic changes and their inheritance (Uller et al., 2015).

Cytosine DNA methylation is a heritable epigenetic modification involving the addition of a methyl (-CH3) group to the fifth carbon of the pyrimidine ring of cytosine nucleotides. This addition is catalyzed by DNA methyltransferases, commonly found in most eukaryotes (Cheng, 1995). In plants, DNA methylation can occur in three sequence contexts: the symmetric CG and CHG contexts, and the asymmetric CHH context, where H represents A, C, or T nucleotides (Law and Jacobsen, 2010). Methylation in these different contexts displays distinct genomic patterning within genes, repeat regions, and transposable elements. While CG methylation is largely concentrated within genes and transposable elements, CHG and CHH methylation contexts are usually associated with repeat regions and transposable elements (Cokus et al., 2008).

One role of DNA methylation is to silence transposable elements, which can become activated during stress conditions (Slotkin and Martienssen, 2007). In some cases, changes in DNA methylation have also been associated with stress-induced gene regulation, such as during phosphate starvation or *Pseudomonas syringae* infection (Dowen et al., 2012; Yong-Villalobos et al., 2015), and may provide the mechanistic basis for memory (Dowen et al., 2012; Kinoshita and Seki, 2014). Despite major progress in dissecting the genetic pathways responsible for establishment and maintenance of context-specific DNA methylation patterns (Stroud et al., 2013), functions of DNA methylation, particularly genic CG methylation, has remained mysterious (Zilberman, 2017).

*MutS Homolog 1* (*MSH1*) is a plant-specific, nuclear-encoded gene that targets its protein to both plastids and mitochondria. Arabidopsis *msh1* mutants display a range of altered phenotypes that include variegation, dwarfing, delayed maturity transition, delayed flowering, and partial male sterility (Xu et al., 2011). The *msh1* mutants display higher tolerance to heat, high light, and drought stress (Shedge et al., 2010; Virdi et al., 2016; Xu et al., 2011), particularly in individuals showing stronger developmental phenotypes. *MSH1* phenotypes are conserved between monocots and eudicots. This conservation is evidenced in the RNAi suppression phenotypes, and the consistent observation that subsequent MSH1-RNAi transgene segregation gives rise to transgenerational *msh1* memory in sorghum, pearl millet, tomato, tobacco and soybean (de la Rosa Santamaria et al., 2014; Raju et al., 2017; Xu et al., 2011; Xu et al., 2012; Yang et al., 2015).

Disruption of *MSH1* causes genome-wide methylome repatterning in both CG and non-CG context (Virdi et al., 2015), along with large-scale changes in gene expression related to abiotic and biotic stress response, phytohormone pathways, circadian rhythm, defense response and signaling (Shao et al., 2017). Arabidopsis *msh1* memory lines show a subset (ca 10%) of the gene expression changes of the T-DNA insertion mutant, with enrichment in circadian rhythm, ABA signaling, and light-response pathways, and with methylome repatterning predominantly in CG context (Sanchez et al., 2018).

In this study, we investigated the stress response behavior of plants following *msh1* developmental reprogramming. We show that *msh1* mutants display a differential response to abiotic and biotic stress, which could be partly explained by transcriptome changes. Epi-lines, deriving from crosses of *msh1* with wild type, showed increased seed yield and higher tolerance to salt, freezing and mild heat stress. Under prolonged cold stress, *msh1* mutants showed increased variation in DNA methylation, particularly in non-CG context, and this increased CHG and CHH methylation pattern variation did not appear to influence the *msh1* crossing-derived vigor phenotype. Taken together, the data imply that developmental phenotypes in the *msh1* mutants are caused by large-scale gene expression changes associated with stress response, along with genome-wide methylome repatterning. Methylome changes in non-CG context were disproportionately affected by cold stress and were hyper-responsive to environmental changes, whereas changes in CG context appeared to be stable and to influence plant phenotype.

## MATERIALS AND METHODS

### Plant growth conditions and PCR genotyping

The genetic background used throughout the study was Arabidopsis Col-0 ecotype. For phenotypic measurements, seeds were sown into plastic pots containing Fafard germination mix with Turface MVP added. After 48-72 hrs of cold stratification at 4 °C in a dark chamber, pots were moved to growth chambers set at 22 °C. The *msh1* T-DNA mutant was obtained from Arabidopsis Biological Resource Center (SAIL_877_F01, stock number CS877617) and genotyped as described previously (Shao et al., 2017). Epi-lines were developed by crossing wild type with *msh1* mutants, some of which had been exposed to cold stress (S), and subsequently self-pollinating filial generations. PCR genotyping as previously described (Shao et al., 2017), was performed on the F_2_ population and only plants with wild type *MSH1/MSH1* were evaluated and forwarded. Yield and stress tests were performed on bulked epi-F_3_ populations.

### Abiotic and biotic stress treatments

All stress treatments were performed on wild type Col-0, *msh1* mutants #9, #12-4 and #12-29, epiF_3_ populations derived from crosses WT x *msh1*-N, WT x *msh1*-VD, WT x *msh1*-N(S), and WT x *msh1*-VD(S) that involved the two phenotypic classes of *msh1* mutants, normal phenotype (N) and variegated dwarf (VD), with and without exposure to stress (S).

Seeds for stress treatments were bleach-sterilized and sown on half-strength MS medium containing 1.5% sucrose and 0.5% MES, pH 5.7, solidified with 4% agar in sterile plastic Petri-plates. For 200mM salt germination tests, 11.7 g of NaCl was added to the growth media before sterilization. After 48-72 hrs of cold stratification in a dark room at 4 °C, plates were moved to Percival growth chambers set at 22 °C and 16/8 light/dark cycle. Germination was scored based on root length of more than 3mm at two weeks after plates were moved to the growth chamber.

For freezing tolerance, two-week-old seedlings were cold acclimatized for one week at 4 °C in 12/12 hrs light/dark photoperiod. Freezing tests were performed as previously described (Barnes et al., 2016), with necessary modifications. Specifically, post-freezing plates were placed in a 4 °C dark chamber for 24 hrs before recovery in control growth conditions for 5-7 days. Survival was scored as plants having fully expanded green rosette leaf after recovery. The *sfr2-3* mutant (Moellering et al., 2010), used as negative control, was a kind gift from Dr. Rebecca Roston.

Two independent *MSH1* epi-lines for each phenotypic class of *msh1* mutant were developed, WT x *msh1*-N1 and WT x *msh1*-N2, created by crossing two independent *msh1* mutants with a normal phenotype (N1, N2), and WT x *msh1*-VD1 and WT x *msh1*-VD2 developed from two *msh1* mutants with a variegated dwarf phenotype (VD1, VD2). Seed yield was measured as total seed weight at maturity. Floral stems of six-week-old plants were tied to a wooden stake and the plant enclosed completely using Arabisifter (Lehle Seeds, SNS-03), making a pouch-like structure in the bottom to collect shattered seeds. All four epi-F_3s_ and wild type were grown in a completely randomized design in a growth chamber at 22 °C or 32 °C, 16/8 hr light/dark cycle. Seeds were carefully harvested from each population (n>18 plants) at maturity. Seeds were dried in a 37 °C chamber for 48-72 hrs before recording seed weights.

*Pseudomonas syringae pv. tomato DC3000* strains were grown for 24 hr at 30 °C on King’s B media (King et al., 1954) with the appropriate antibiotics, and resuspended to an OD_600_ of 0.2 (2 x 10^8^ cells ml^−1^) in 10 mM MgCl_2_. The resuspended culture was sprayed uniformly on upper and lower surfaces of fully expanded leaves of 4-week-old wild type, *msh1* mutant, and *msh1*-derived epi-lines using a jet-spray bottle. Treated plants were well-watered and kept in a dark room for five days, followed by five to seven days in a growth chamber at 22 °C and 16/8 hr light/dark cycle before scoring for survival.

### RNA extraction and sequencing analysis

Four-week-old plants grown in 22 °C were transferred to a growth chamber set at 10 °C, 150 μE m^−2^ s^−1^, for 30 days. Tissue from four fully expanded rosette leaves was sampled before and after 10 °C transfer with three replicates per group. For each sample, frozen tissue was ground and total RNA extracted using a standard TRIzol reagent protocol. RNA samples were then treated with DNaseI (Qiagen catalog #79254). Qiagen RNeasy Plant Mini Kit (Qiagen catalog #74904) was used to purify total RNA samples prior to RNA sequencing (RNAseq). Poly(A)-enriched RNAseq was performed by Beijing Genomics Institute (BGI), generating at least 59.6 M paired-end, 100-bp reads per sample. Reads were trimmed and aligned to the Arabidopsis TAIR10 reference genome sequence with annotation from Araport11 PreRelease3 using TopHat2 (Kim et al., 2013). The DESeq2 method (Love et al., 2014) was used to identify differentially expressed genes (cutoff of FDR < 0.05, |log2(fold change)| ≥ 1, and mean FPKM ≥ 1). Gene Ontology (GO) enrichment analysis was performed using the DAVID database (Huang et al., 2009). GO terms with p-value < 0.05 after Benjamini-Hochberg (Benjamini and Hochberg, 1995) correction for multiple testing were considered statistically significant in each comparison.

For transposable element (TE) family expression analysis, reads were aligned using the STAR 2-pass method (Dobin et al., 2013), allowing up to 100 multi-mapped locations as per the recommendation of TEtranscripts (Jin et al., 2015). Quantification and testing for differential expression of TEs were performed using TEtranscripts with the developer-provided Arabidopsis TE family annotation.

### Cold stress methylome analysis

To obtain whole-genome bisulfite sequencing data for the cold stress experiment, plants were grown in a controlled growth chamber set to 10 °C, 150 μE m^−2^ s^−1^ or 500 μE m^−2^ s^−1^ and 12/12 day/night photoperiod for 21 days, beginning from germination, then moved to recovery at 22 °C, 250 μE m^−2^ s^−1^ for 18 days before sampling. Control plants were grown continuously at 22 °C, 250 μE m^−2^ s^−1^ from sowing, and sampled upon reaching a similar developmental stage as cold-stress recovered plants. Four fully-expanded rosette leaves from each individual plant were harvested and DNA extracted as previously described (Li and Chory, 1998), with two replicates per group. Library generation and bisulfite-sequencing were performed by BGI on a Hiseq2000. Reads were aligned to the TAIR10 reference genome using Bismark (Krueger and Andrews, 2011) with default mismatch parameters. Due to the potential for artifacts, cytosines of CCC context were excluded from CHH analysis.

The R package *methylKit* 1.1.8 (Akalin et al., 2012) was used to call DMRs, based on 100 bp non-overlapping windows, separately for CG, CHG and CHH contexts. Only cytosine base positions with ≥ 3 reads were retained for analysis, and normalized methylation counts for each cytosine were used based on coverage. Windows with ≥ 5 cytosines (of the given context) were considered for analysis, to rule out low information regions. The principal component analysis was performed using the *PCASamples* function. Subsequent comparison between treatment (cold or control) and genotype (*msh1* T-DNA or wild type) combinations were performed by logistic regression with methylKit. DMRs for each context were identified based on a methylation difference of at least 10% absolute value and a q-value < 0.05, then clustered using Ward’s method (Ward Jr, 1963). For CG context, genes overlapping DMRs within each cluster were identified and subjected to GO enrichment analysis using the DAVID database (Huang et al., 2009). For CHG and CHH contexts, TE’s overlapping DMRs within each cluster were identified and tested for enrichment of TE families and superfamilies (annotated by TAIR10) using the hypergeometric test (FDR < 0.01).

## RESULTS

### The *msh1* mutant shows variable abiotic and biotic stress tolerance

Previous studies have shown that *msh1* mutants are more tolerant to drought, high light, and heat stress (Shedge et al., 2010; Virdi et al., 2016; Xu et al., 2011). We tested for other abiotic stress effects, focusing first on salt and freezing temperature. Seeds of *msh1* mutant and wild type were grown on plates with half-strength MS media and 200 mM NaCl. The 200 mM NaCl concentration is highly selective for germination tests in Col-0 (Wibowo et al., 2016). Germination was scored based on root length of greater than 3mm at two weeks after sowing, assessed in three independent experiments. Only 32% percent of wild type seeds germinated on 200mM NaCl plates. Two *msh1* mutants, #9 and #12-29, showed significantly higher germination than wild type (p-value 1.25e-10 and 0.000128 respectively), while *msh1*#12-4 did not show significant difference (p-value 0.147672). These results suggest higher salinity tolerance in *msh1* mutants, with variation in mutant sub-populations (Fig 1A). This result is consistent with gene expression data from *msh1* mutants (Shao et al., 2017), which show differential expression for 493 (Dataset S1) of the 1667 salt stress-responsive genes identified through comparative microarrays (Sham et al., 2015).

**Figure 1.**
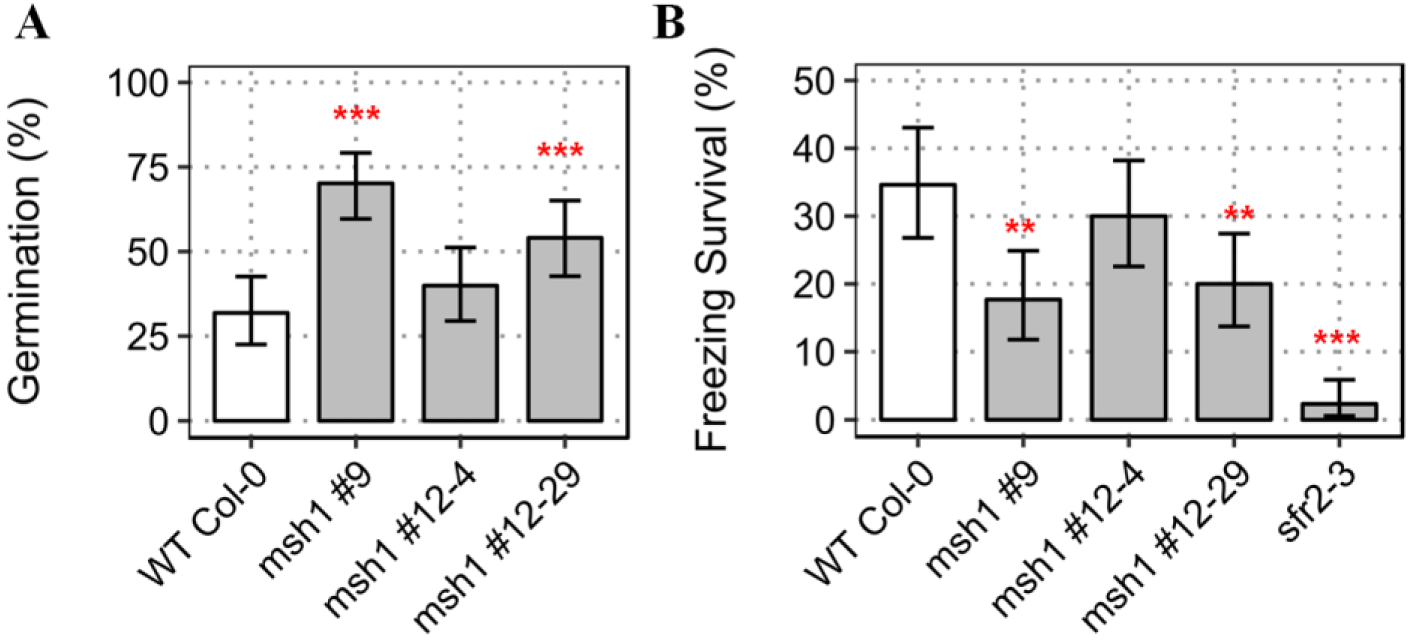
Abiotic stress tests in Arabidopsis *msh1* mutants. **A)** Percent germination rate of wild type Col-0 and, *msh1* mutants #9, #12-4, and #12-29 at 200mM NaCl-supplemented growth media, scored after two weeks post sowing [n=100 plants each; error bars represent standard error of means (SEM)]. **B)** Proportion of recovered (survived) plants seven days post freezing treatment at -10 °C. *sfr2-3* was used as negative control for freezing tolerance (n=130, error bars represent SEM). Significance at '***' 0.001 '**' 0.01 '*' 0.05 '.' 0.1

To examine whether or not *msh1* mutants also showed tolerance to freezing temperatures, two-week-old seedlings of wild type and *msh1* T-DNA mutants were cold acclimatized for a week before exposure to -2 °C for 4 hrs, followed by nucleation and -10 °C for 12 hrs. Survival was scored as the presence of green rosette leaves one week after recovery under normal growth conditions (22°C, 16/8 light/dark cycle). Surprisingly, the survival rate of *msh1* mutants was lower than that of wild type (Fig 1B), indicating that the mutants are not tolerant to all stresses. Under our experimental conditions, 34.5% wild type survived -10 °C. The *msh1* mutants #9 and #12-29 showed significantly higher susceptibility to freezing temperatures (p-value 0.00222 and 0.00881respectively), while *msh1* mutant #12-4 was not significantly different from wild type (p-value 0.42651). From a set of 590 differentially expressed genes correlated with acclimated and non-acclimated freezing tolerance (Hannah et al., 2006), only 64 were altered in expression in the *msh1* variegated dwarf mutant (Dataset S2).

Because *msh1* mutants have increased tolerance to abiotic stresses like drought, heat, high light, and salt, we tested whether or not they were likewise more resistant to biotic stress. We challenged *msh1* mutants with the gram-negative bacterial pathogen *Pseudomonas syringae* pv. *tomato* DC3000, which causes bacterial speck disease in tomato and is pathogenic to Arabidopsis. The *msh1* mutants showed susceptibility to the bacterial pathogen. While 87.5% of wild type plants survived the stress, *msh1* mutant sub-populations #12-29 and #12-4 showed significantly higher susceptibility (Fig S1A: p-value 0.00633 and 0.08136 respectively). Within one population, *msh1* #9, plants with variegation and dwarfing showed significantly higher susceptibility (p-value 4.35e-05 and 7.15e-07 respectively) to *P. syringae* than *msh1* mutants with a mild phenotype (Fig S1B: p-value 0.327). Thus, *msh1* mutants are susceptible to biotic stress despite markedly increased expression of biotic stress-responsive pathways in *msh1* mutants (Shao et al., 2017), and the biotic stress response appears related to the severity of the *msh1* phenotype.

The *msh1* mutant and derived *msh1* memory lines are considered to represent two distinct epigenetic states of *msh1* effect, differing in phenotype, methylome and gene expression profiles (Sanchez et al., 2018), with a third state emerging from crosses of the *msh1* mutant (or memory line) to wild type. This third state is characterized by markedly enhanced growth vigor (Virdi et al. 2015). To investigate the inheritance of these stress responses following *msh1* crossing, seed germination rate in 200mM NaCl concentration and survival of seedlings at -10 °C freezing temperatures were assayed in three independent experiments for *msh1*-derived epi-lines in the F_3_ generation, with wild type as a control. Epi-lines were created by crossing wild type Col-0 with *msh1* mutants as pollen donor, and self-pollinating filial generations to obtain epi-F_3_ bulks (see methods). When seeds were germinated on plates with 200mM NaCl, WT x *msh1*-N and WT x *msh1*-VD showed significantly higher germination rate (p-value 7.15e-14 and 3.20e-12 respectively) than wild type (Fig S2A). This result was consistent with the parental *msh1* mutant, which showed a similar increase in salt tolerance (Fig 1A). However, epiF_3_ population WT x *msh1*-VD also showed higher tolerance to freezing (Fig S2B: p-value 0.014546), where *msh1* mutant showed greater susceptibility. These observations are consistent with the expectation that *msh1* x wild type crosses produce a different epigenetic state, thus resulting in distinctive phenotypes. Crossing the *msh1* mutant may alter circadian clock regulation of freezing stress response.

To evaluate the response of progeny from crossing under less severe, non-lethal stress, we subjected epi-F_3_ plants to mild heat stress and measured total seed weight at harvest. For this experiment, four epi-lines and wild type were grown in growth chambers under control (22 °C) or mild heat stress (32 °C) throughout the plant life cycle. Epi-lines showed 9.7% to 19.6 % increase in seed yield compared to wild type in control conditions (Fig 2A). Three of the epi-lines also performed significantly better than wild type under mild heat stress, showing 9.5% to 16.5% increase in yield (Fig 2A). The lower yield penalty under mild heat stress in the three epi-lines, coupled with the enhanced salt and cold tolerance, provides an indicator of greater yield stability and lower environmental effects on the *MSH1* growth-enhanced phenotype (Fig 2B). These results resemble the higher yield stability observed in soybean *MSH1* epi-lines grown across four different locations in Nebraska (Raju et al., 2017).

**Figure 2.**
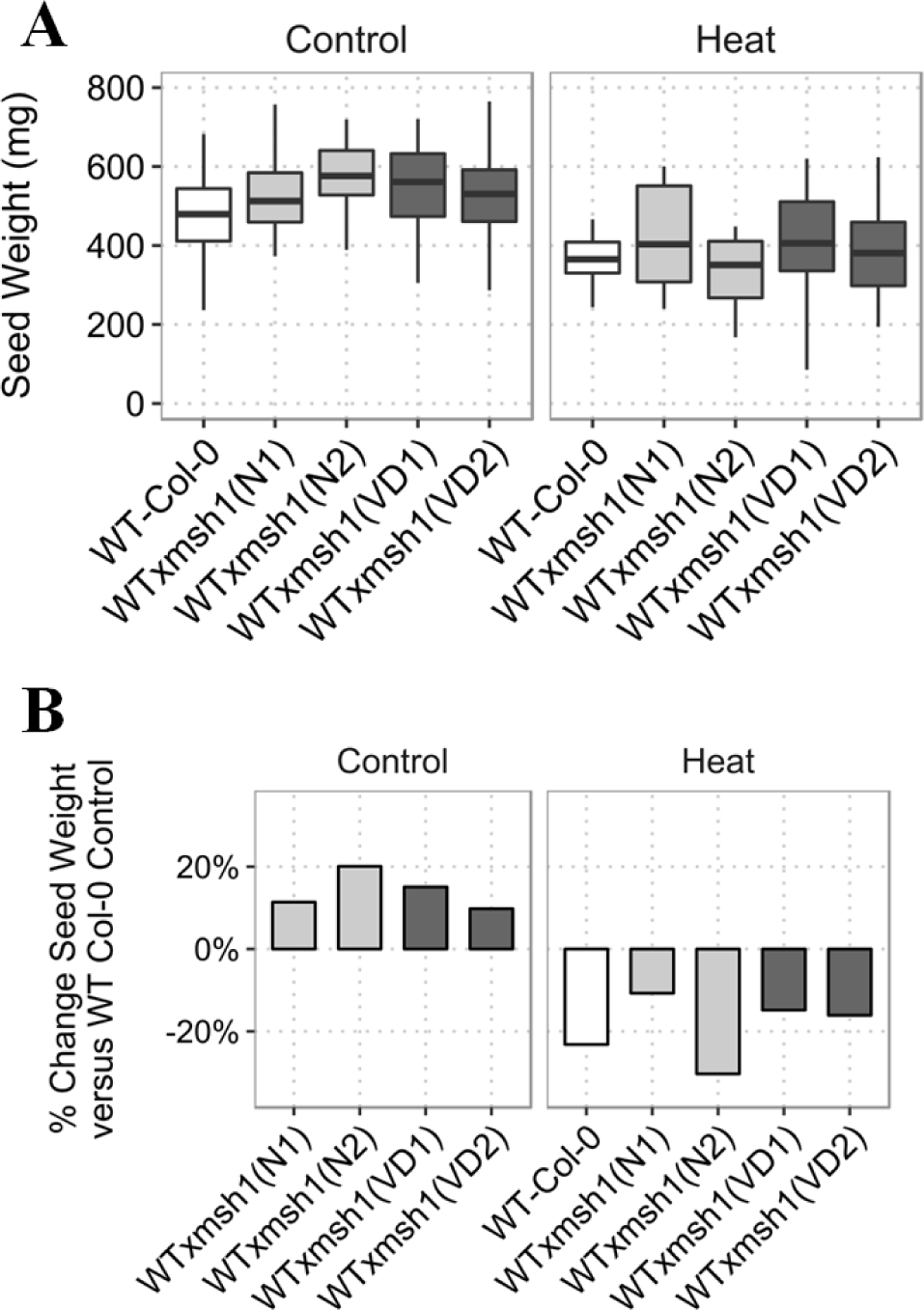
Total yield measurements for *msh-* derived epi-lines compared to wild type Col-0. **A)** Whisker plots showing differences in seed yield between wild type and *msh1* epi-lines under control (22 °C) and mild heat-stress (32 °C) growth conditions [Control (n=36), heat stress (n=18)]. **B)** Percent change in seed weight for epi-lines under control and heat stress condition compared to seed weight of wild type under control growth conditions.

### The methylome of *msh1* is hyper-responsive to cold stress with disproportionately higher CHH hypomethylation

Transcriptome studies of *msh1* showed clear enrichment of biotic and abiotic stress response genes, including response to cold. Despite changes in cold-responsive transcription factors (Shao et al., 2017), *msh1* mutants showed susceptibility to freezing temperatures. These observations led us to test whether *msh1* mutants would show differential methylome and transcriptome response to low-temperature stress.

To evaluate the extent of DNA methylation changes related to long-term cold stress, *msh1* mutants and wild type plants were grown at 10 °C for 18 days under 12/12 light/dark cycle, then allowed to recover at 22 °C for 18 days before sampling for DNA extraction. Plants were allowed to recover prior to sampling for two reasons: Plant growth was slower under cold stress, complicating the collection of sufficient tissue for methylome sequencing. In addition, we wanted to avoid transient methylation changes present during plant exposure to cold treatment.

To facilitate comparison of each region between different genotype and treatment combinations, methylome analysis was performed using fixed 100-bp non-overlapping windows. Principal component analysis plots from the first two principal components using the upper 0.9 quantile of variable windows showed CG methylation separating by genotype between wild type and *msh1*mutants, with or without stress (Fig 3A). These observations are consistent with studies of the *msh1* memory lines, where CG methylation is predominant in association with a memory phenotype (Sanchez et al., 2018). CHG methylation showed a similar pattern, although cold-stressed samples were discriminated from control samples in *msh1* mutants more than in wild type (Fig 3A). Notably, CHH methylation showed the greatest degree of discrimination for the cold stress treatment, predominantly in *msh1* mutants (Fig 3A). Together, these results indicate that cold stress influences DNA methylation in all methylation contexts, but there is evidence of interaction with the *msh1* background, amplifying the effect in CHH context.

**Figure 3.**
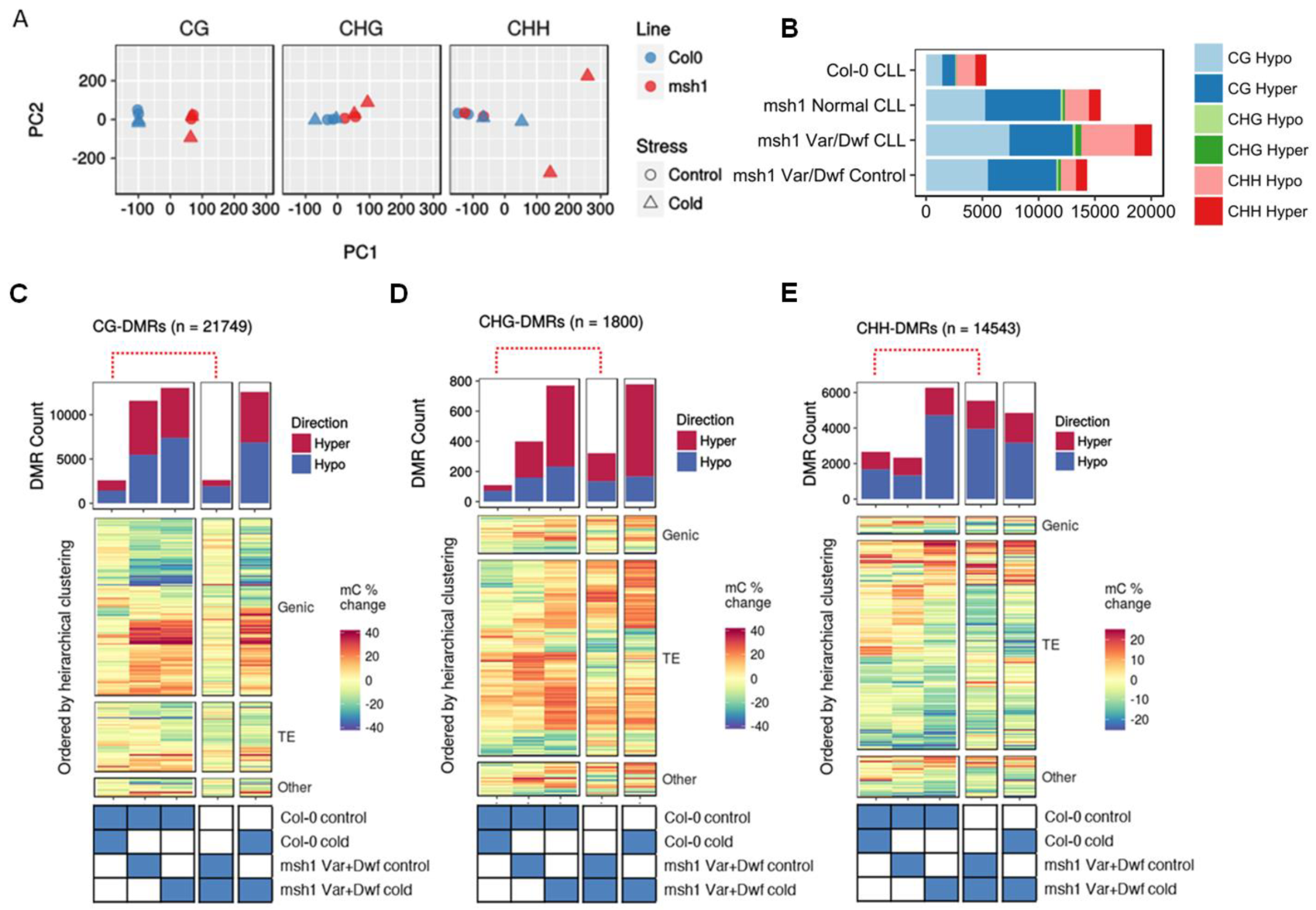
Methylome changes in *msh1* mutants and wild type Col-0 from long-term cold stress. **A)** Principal component analysis (PCA) plots for methylation levels within 100-bp windows separated for nucleotide context; CG, CHG, and CHH (H represents A, C, or T). **B)** Graph of total DMR numbers in each comparison, showing hyper and hypomethylation in all three contexts. **(C-E)** DMR counts and hierarchical clustering of all pair-wise comparisons, for CG **(C)**, CHG **(D)**, and CHH **(E)** contexts. Red dotted lines highlight *msh1* cold response relative to wild type under cold stress.

### Genome-wide distribution of DMRs in wild type and *msh1* mutants in response to cold stress

We investigated the number and genomic distribution of differentially methylated regions (DMRs). DMR calling was based on logistic regression over 100 bp non-overlapping window, using a threshold of more than 10% absolute change in methylation level in each cytosine context. The resulting number of DMRs (Table 1, Fig 3B) confirmed trends observed by principal component analysis (Fig 3A). As expected, CG-DMRs mostly occurred over genes and were relatively few between cold and control treatments in wild type or *msh1* while comparing any *msh1*group to wild type (Fig S3A). We found 11,579 CG-DMRs, 399 CHG-DMRs, and 2332 CHH-DMRs when comparing *msh1* to wild type under control conditions. Almost equal numbers of DMRs were hyper or hypomethylated in symmetric methylation context, while in CHH context there were 30% more hypomethylated DMRs in *msh1* (Table 1, Fig 3B). We also detected 2592 CG-DMRs, 109 CHG-DMRs and 2658 CHH-DMRs induced by cold stress alone in the wild type. The magnitude of CG changes in the *msh1* mutant was 4.35 times higher than CG changes induced by cold stress in wild type, while the magnitude of CHH changes was not significantly different. This implies that cold stress predominantly affects CHH methylation, more than CG and CHG methylation, consistent with previous reports of methylome behavior under low temperature (Dubin et al., 2015).

**Table 1:** Number of DMRs in all three cytosine contexts across multiple comparisons.

| Comparison | Direction | CG | CHG | CHH |
| --- | --- | --- | --- | --- |
| *msh1* vs WT | Hyper | 6085 | 240 | 996 |
|  | Hypo | 5494 | 159 | 1336 |
| *msh1*(S) vs *msh1* | Hyper | 700 | 186 | 1574 |
|  | Hypo | 1926 | 135 | 3965 |
| WT(S) vs WT | Hyper | 1154 | 37 | 977 |
|  | Hypo | 1438 | 72 | 1681 |
| *msh1*(S) vs WT | Hyper | 5626 | 538 | 1548 |
|  | Hypo | 7400 | 232 | 4723 |
| *msh1*(S) vs WT(S) | Hyper | 5707 | 611 | 1675 |
|  | Hypo | 6829 | 167 | 3176 |

We examined whether *msh1* background affects methylation changes upon cold stress. We found 2626 CG-DMRs, 321 CHG-DMRs and 5539 CHH-DMRs between stressed and unstressed *msh1* mutant (Table 1, Fig 3B). Thus, CHH methylation, primarily over transposable elements (Fig S3A), showed the greatest effect of cold treatment within the *msh1* background, consistent with separation seen in the PCA plot (Fig 3A). Although CHH methylation is affected by cold stress in wild type, CHH DMRs in cold-stressed *msh1* are twice as abundant as in similar wild type comparisons. Whereas CG DMR patterns were nearly identical for cold-stressed and control *msh1* mutants when compared to wild type, CHG and CHH DMRs showed a clear distinction in patterns, with several loci switching between hyper and hypomethylation (Fig 3C-E). These results indicate an interaction between the *msh1*effect and cold stress, such that non-CG methylation patterns are disproportionately affected.

### Non-CG methylome changes in association with transposable elements

To investigate the genomic distribution of non-CG changes in response to stress, we clustered non-CG DMRs and looked for enrichment of TE superfamilies in these clusters. Both CHG- and CHH-DMRs formed 4 clusters each (Dataset S3). While all four clusters in CHG-DMRs showed enrichment for DNA/En-spm, LTR/COPIA, and LTR/Gypsy elements, clusters three (hyper) and one (hypo), which showed similar trends in all comparisons, were also enriched in LINE/L1 elements. In CHH-DMR clusters, DNA/MuDR elements were enriched in all clusters. Cluster one, which contained the most DMRs and hypomethylation in all three comparisons (wild type-stressed vs wild type, *msh1* vs wild type, and *msh1*-stressed vs wild type) showed enrichment for LINE/L1, LTR/COPIA, and LTR/Gypsy elements. Clusters three and four, which showed hypermethylation in *msh1*-stressed versus wild type, showed over-representation of DNA/Mariner and RC/Helitron elements.

Genomic distribution of DMRs matched with known behaviors within each cytosine context. CG-DMRs between *msh1* mutants and wild type were distributed evenly across the chromosome (Fig S4B, C, E), while CG DMRs from cold stress were primarily limited to heterochromatin (Fig S4A, D). CHG-DMRs and CHH-DMRs were mainly in heterochromatic regions for both comparisons. This finding is consistent with previous reports of cold stress methylome changes showing heterochromatin bias (Dubin et al., 2015). We examined expression changes in genes related to DNA methylation machinery. Interestingly, CHROMO METHYLTRANSFERASE 3 (CMT3) and DECREASE IN DNA METHYLATION 1 (DDM1) expression were down-regulated in cold-stressed *msh1* mutants compared to unstressed mutants and wild type (Fig S5). Since *cmt3* and *ddm1* mutants are known to increase heterochromatic TE de-repression, these observations appear consistent with CHH hypomethylation of heterochromatin in the interaction of *msh1* effect and low-temperature stress.

### Transcriptome response of Arabidopsis *msh1* mutants under chronic cold stress

We evaluated the effect of cold stress on the transcriptome of *msh1* mutants. Wild type Col-0 and *msh1* plants were grown at 22 °C for four weeks before leaf tissue was collected (control group), or grown at 10 °C for an additional 30 days before sampling tissues for RNA extraction (cold-stressed group). Transcriptome analysis showed cold stress to be the largest contributor to transcriptional changes within the experimental groups, evident from the groups formed in PCA plotting with normalized log values of gene expression (Fig 4A). Although the magnitude of gene expression change was lower than transcriptome change in our earlier report (Shao et al., 2017), similar pathways were modulated in both *msh1* mutants with or without severe phenotype, including defense, jasmonic acid, abiotic stress response, photosynthesis and oxidative stress (Dataset S6). Technical differences, like differential developmental staging and changes in circadian phase (Hsu and Harmer, 2012), might explain the differences in the magnitude of transcriptome changes. Pathways affected in *msh1* appear to be induced by cold alone in wild type, suggesting that *msh1* mutants have stress response pathways activated in the absence of any environmental cues. Response to abiotic stress (cold, salt, light, and wounding) and biotic stress (response to chitin and jasmonic acid) are activated as a cold stress response in wild type and are also activated in *msh1* (Fig S6A). Defense response, jasmonic acid-mediated signaling, and photosynthesis-related genes were specifically enriched in *msh1* (Fig S6A: Dataset S6). Taken together, these results suggest that unlike methylome, transcriptome changes do not show increased plasticity in *msh1* mutants under cold stress.

**Figure 4.**
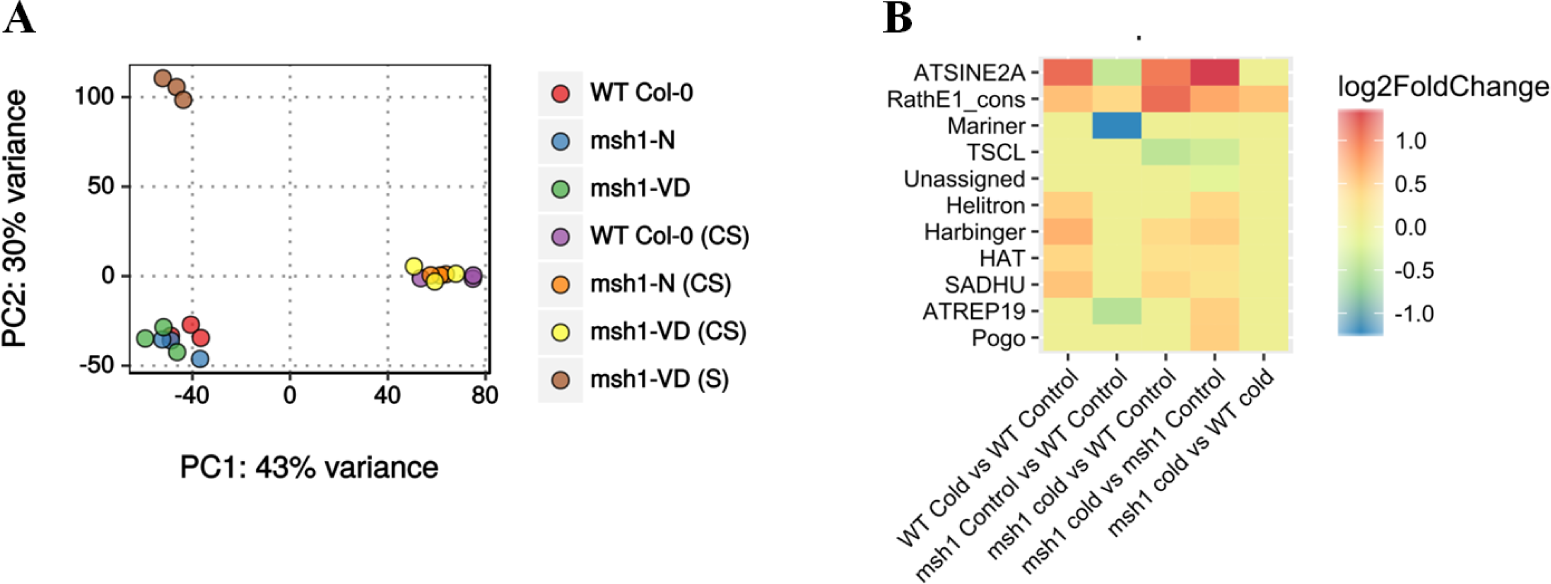
Transcriptome changes in *msh1* mutants before and after chronic cold stress. **A)** PCA plot from normalized log values of gene expression from wild type and *msh1* mutants under control or cold stress. **B)** Heat map showing differential expression of transposable element (TE) super families from each corresponding comparison within cold stress experiment.

We also looked into differences in transposable element expression using TEtranscripts (Jin, et al. 2015). Mariner, ATREP19, and SINE TE superfamilies were significantly down-regulated in *msh1* mutants compared to wild type, while Rath elements were up-regulated in all comparisons (Fig 4B). In contrast, SINE elements showed clear cold-stress induced up-regulation. Similarly, Helitron, Harbinger, HAT, and SADHU elements were up-regulated in wild type under cold stress (Fig 4B). At the family level, ATCOPIA28 and ATCOPIA31A showed clear stress-induced up-regulation, while VANDAL5A, ATREP3, ATCOPIA44, ATCOPIA 78, ATCOPIA 93, and ATMU1 showed down-regulation in *msh1* mutants (Fig S6B).

### *MSH1*-induced CG methylation changes are associated with enhanced growth in progeny from *msh1* crossing

To evaluate the extent to which non-CG methylome divergences affect the *msh1* crossing-derived enhanced growth phenotype in Arabidopsis (Virdi et al., 2015), we investigated the epi-lines from *msh1* mutants with or without cold stress (see methods). We assayed rosette diameter, days to flowering, and total seed weight from F_2_ and selected F_2:3_ lines under control growth conditions. The F_2_ population WT x *msh1*-N(S) showed higher mean rosette diameter compared to wild type, measured at six weeks after sowing (Wilcox test, padj 0.004, Fig 5A). This population flowered an average of two days earlier (Wilcox test, padj 0.002, Fig S7). Both WT x WT(S) and WT x *msh1*-N populations showed smaller rosette diameter compared to wild type (Fig 5A). Mean seed yield, measured in milligrams, for WT x *msh1*-VD(S) and WT x *msh1*-VD was significantly higher than wild type Col-0 (Wilcox test, padj 0.015, Fig 5B), whereas no significant difference was found between WT x *msh1*-VD(S) and WT x *msh1*-VD (t-test, p-value 0.80). These results show that for epiF_2s_, WT x *msh1*-VD(S) and WT x *msh1*-VD showed 20% and 17.9% increase in yield compared to wild type, but stressing the *msh1* mutant prior to crossing does not have a significant effect on yield. We also noticed that while WT x *msh1*-VD and WT x *msh1*-VD(S) showed increases in seed yield, WT x *msh1*-N(S) showed higher rosette diameter compared to wild type, suggesting the possibility of selection for separate traits of vegetative biomass heterosis and seed yield heterosis in *msh1* derived epi-lines.

**Figure 5.**
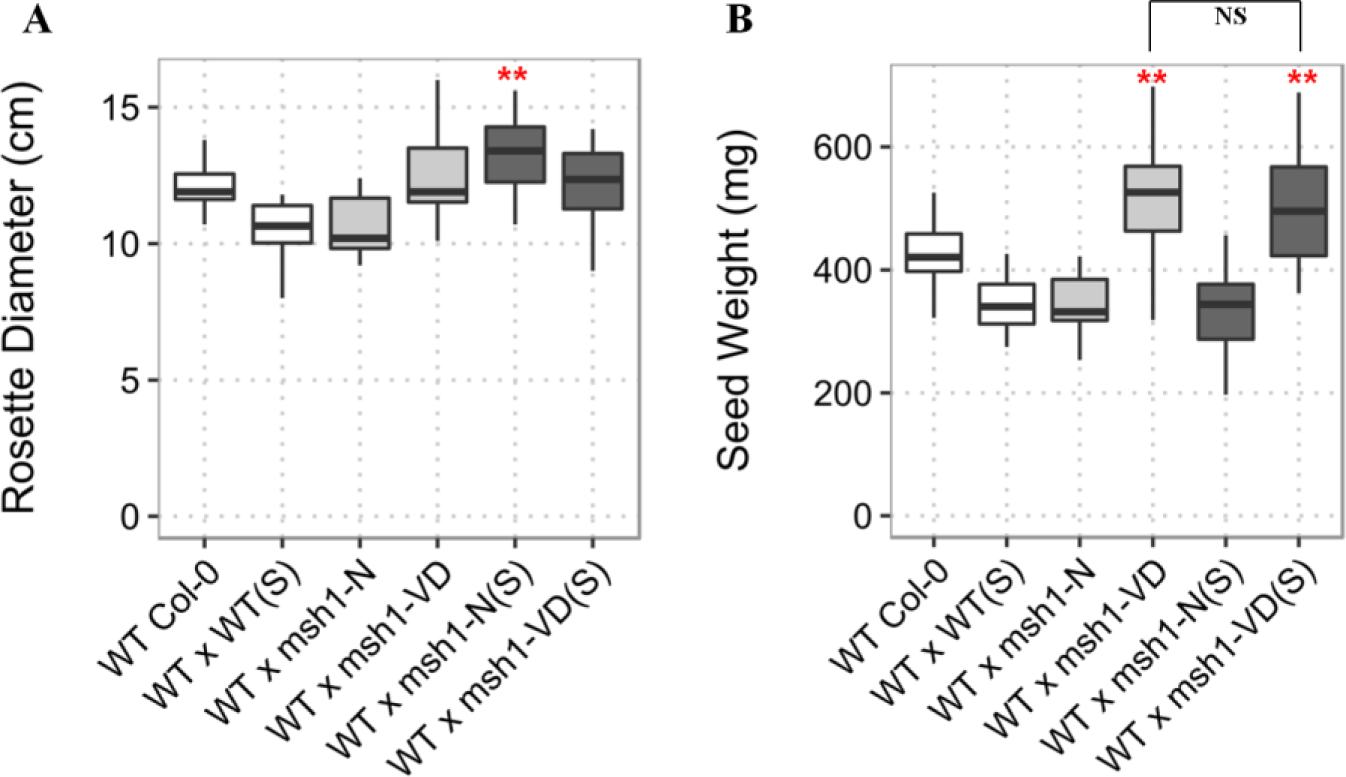
Variation in rosette diameter and seed weight in *MSH1* epi-F_2_ population. A)Whisker plot showing variation in rosette diameter measured at 6 weeks after sowing (n= 18). B)Whisker plot showing variation in total seed weight measured carefully after bagging the plants with Arabisifter (Lehle Seeds, SNS-03). Epi-F_2_ populations were developed from cold stressed (S) and unstressed *msh1* mutants with (VD) or without (N) phenotype. F_2_ plants were selected after genotyping for *MSH1/MSH1* wild type allele. Rosette diameter and seed weight were measured from the same set of F_2_ plants. Significance at "***" 0.001 "**" 0.01 "*" 0.05 "." 0.1

We evaluated total seed weight for F_2:3_ lines following selection of the upper 20% for seed weight in each F_2_ population under control conditions. Although the selection was performed on seed weight, F_2:3_ epi-lines 3C2, derived from WT x *msh1*-VD(S), and 4C2, derived from WT x *msh1*-VD, showed significantly higher rosette diameter (Wilcox test, padj = 0.018, Fig 6A) compared to wild type. Both epi-lines also showed significantly higher seed weight compared to average wild type (t-test, p-value = 0.0008 and 0.043 respectively, Fig 6B), reflecting a response to selection. Similar to F_2_ results, there was no significant difference in seed weight between 3C2 and 4C2 (t-test, p-value = 0.203), confirming that stress treatment of *msh1* mutants does not have an effect on *msh1*-derived growth enhancement in progeny from crosses.

**Figure 6.**
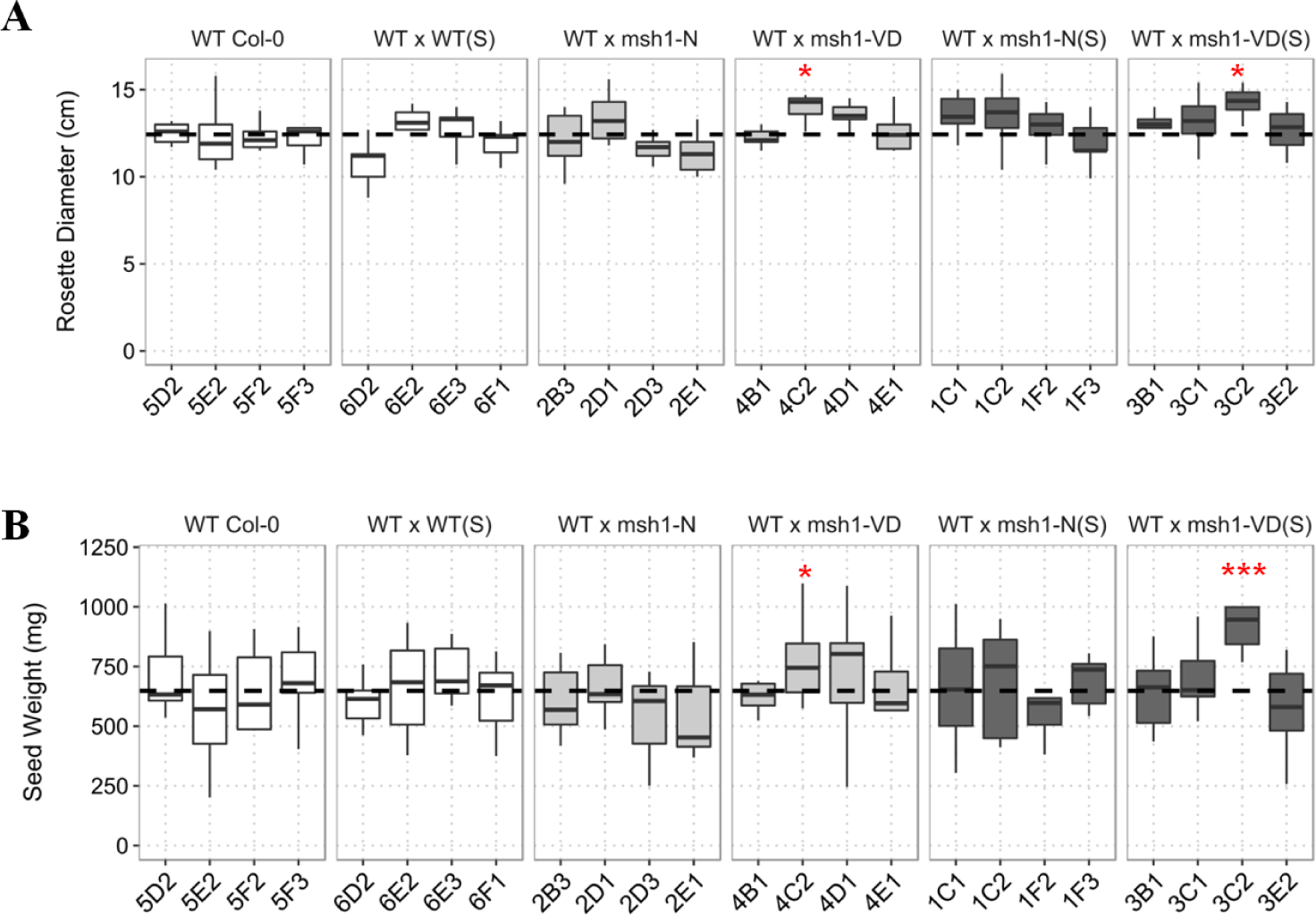
Rosette diameter and total seed weight in selected epiF_2:3_. **A)** Seed weight measurements from top 20% selection in each epi-F_2_ population, including four sub-lines for wild type and WT x WT(S). Seed weight was measured in milligrams dried seeds collected from plants bagged with Arabisifter (Lehle Seeds, SNS-03). Black dotted line represents wild type average (n=9 plants each). **B)** Rosette diameter measured from the same plants at six weeks post sowing. Significance at "***" 0.001 "**" 0.01 "*" 0.05 "." 0.1

We subsequently tested whether or not observed methylome changes had an effect on stress adaptation of derived epi-lines. Surprisingly, epi-lines derived from the cold-stressed *msh1* mutant as parent showed a different response to salt and freezing stress. Whereas WT x *msh1*-VD(S) and WT x *msh1*-N(S) epi-F_3_ populations were significantly more tolerant to salt stress (p-value 0.000914 and 0.001803 respectively), although lesser in magnitude to comparable populations from unstressed (Fig 7A), they were not significantly different from wild type in their response to freezing stress (p-value 0.867143 and 0.903767 respectively, Fig 7B). These results suggest that an interaction exists between *msh1* effect and cold stress effects. Taken together, data indicate that stressing *msh1* mutants triggers a disproportionate increase in non-CG methylation, but these changes do not affect *msh1*-derived growth vigor, and can negatively affect stress adaptation in *msh1*-derived epi-lines.

**Figure 7.**
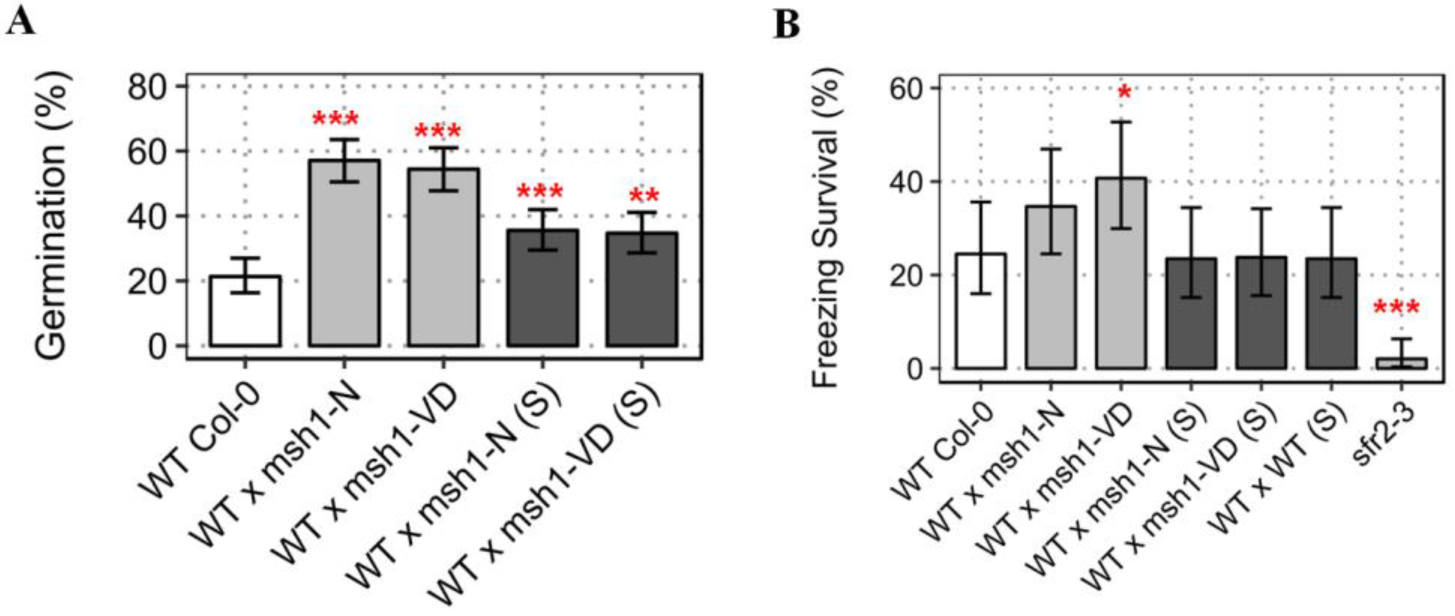
Abiotic stress tolerance in epi-F_3s_ derived from cold-stressed (S) and unstressed *msh1* mutants. **A)** Percent germination of wild type Col-0 and epi-F_3_ bulks; WT x *msh1*-N, WT x *msh1*-VD, WT x *msh1*-N(S), WT x *msh1*-VD(S) at 200mM NaCl-supplemented growth media. Each bar represents three replicates (n=225) and error bars show SEM. **B)** Percent survival of wild type and epi-F_3s_, after 12 hrs at -10 °C. *sfr2-3* was used as negative control. Survival was scored after one week recovery from three replicates (n=100), error bars represent SEM. Significance at '***' 0.001 '**' 0.01 '*' 0.05 '.' 0.1

## DISCUSSION

Previous studies have shown *msh1* mutants to be more tolerant to high light, drought and heat stress (Shedge et al., 2010; Virdi et al., 2016; Xu et al., 2011), consistent with enrichment for abiotic stress response genes (Shao et al., 2017). While we saw increased tolerance for salt stress, *msh1* mutants showed lower survival rate at freezing temperature and in response to the bacterial pathogen *P. syringae*. The seeming incongruity between activation of multiple stress pathways and susceptibility to freezing temperatures may be due to specific mechanisms underlying freezing tolerance in plants, which include plastid membrane remodeling (Moellering et al., 2010). Indeed, low frequency, localized plastid genome changes are reported in *msh1* mutants, along with a reduction in the number of plastids per cell and dramatically altered thylakoid membrane structure in a portion of the organelle population (Xu et al., 2011). Also, loss of *MSH1* might affect the functions of its putative protein interactors, such as the PsbP family protein PPD3 (Virdi et al., 2016), which could further impact the plastid. Alternatively, *msh1* mutants may be unable to mount an appropriate response to freezing stress due to desynchronization of the circadian clock (Shao et al., 2017), which influences freezing tolerance (Maibam et al., 2013). Freezing tolerance is impaired in *cca1-11/lhy-21* double mutants, and *gi-3* mutants are susceptible to freezing stress due to impaired sugar metabolism (Cao et al., 2005; Dong et al., 2011). Also, CBF1 and CBF3 genes, which are positive regulators of cold acclimatization (Novillo et al., 2007), are down-regulated in *msh1* mutants (Shao et al., 2017).

A recent study has suggested that miR163 is a negative regulator of defense response to *P. syringae* in Arabidopsis (Chow and Ng, 2017). Interestingly, *msh1* mutants with variegation and dwarfing showed up-regulation of miR163, while *msh1* mutants with subtle mutant phenotype did not show any changes (Shao et al., 2017). The up-regulation of miR163 in *msh1* mutants with pronounced phenotype corresponds well with their susceptibility to the bacterial pathogen, while mutants with mild *msh1* phenotype show survival rates similar to wild type (Fig S1B). Therefore, one possible explanation for observed stress responses is that organellar changes and/or modulation of key regulatory genes might affect particular stress response, while the vast majority of transcriptional changes may comprise a compensatory response that does not affect the phenotypic outcome.

Whole-genome bisulfite sequencing of *msh1* mutants previously revealed numerous changes in DNA methylation over both genic regions and transposable elements (Virdi et al., 2015), raising the possibility of epigenetic feedback as a response to *MSH1* loss, and heritable methylation changes at stress-responsive loci (Kinoshita and Seki, 2014). Enhanced tolerance to salt stress in epi-lines developed by crossing wild type Col-0 with *msh1* mutants supports the heritability of methylation changes at stress-responsive loci. The derived epi-lines also showed tolerance to freezing stress, despite the parental *msh1* mutant showing susceptibility to freezing temperatures. It is possible that circadian regulation may resynchronize following the crossing of *msh1* mutants with wild type, which is known to influence freezing tolerance in plants (Maibam et al., 2013). In comparable soybean epi-F_4_ lines, circadian genes *GI* and *PRR3/5/7* were up-regulated (Raju et al., 2017), suggesting that modulation of circadian regulators follows crosses with *msh1*.

Derived epi-lines have been shown to display higher yield stability through reduced epitype-by-environment effect in soybean multi-location field trials (Raju et al., 2017). In Arabidopsis, we likewise observed a lower yield penalty under mild heat stress in epi-lines compared to wild type (Fig 2B), implying higher buffering across environments. These observation invite more detailed investigation of the link between *msh1* derived epigenetic variation and decreased environment interaction in derived epi-lines.

Long-term cold stress disproportionately affects *msh1* mutants, which show an amplified CHH hypomethylation response primarily in the heterochromatic region. It is notable that epigenetic changes reported in *msh1* mutants under cold stress mainly involve non-CG methylation at TE sites, predominantly retroelements known to be affected by stress (Wessler, 1996). We also see down-regulation of *CMT3* and *DDM1* in cold-stressed *msh1* mutant compared to wild type and unstressed *msh1* (Fig S5), which is implicated in heterochromatic TE derepression.

The *msh1* mutants showed significant differences in expression of TE superfamilies. Differentially expressed TEs belonged to Rath elements, SINEs, and Mariner superfamilies known to contain shorter TEs, on average (Lewsey et al., 2016), that are usually methylated by the DRM1/2 pathway (Stroud et al., 2014; Stroud et al., 2013). Mariner TE sequences are significantly underrepresented in exons and are often absent in GC-rich genic regions of the genome (Lockton and Gaut, 2009).

Transcriptome studies showed that stress was consistently the major contributor to gene expression changes in wild type and *msh1* mutants. This is expected since changes in CHG and CHH methylation in Arabidopsis are concentrated around the pericentromere, while CG changes are distributed throughout the genome, and non-CG methylation changes are unlikely to direct gene expression changes.

A recent study involving multiple ecotypes in Arabidopsis has shown CHH hypomethylation from lower temperatures, with much of the temperature variation in CHH methylation due to components of the RdDM pathway (Dubin et al., 2015). Reports of chromatin changes and epigenetic features of stress memory in plants and observations that some epigenetic mutants have altered stress responses support the argument that these changes may have biological roles (Probst and Mittelsten Scheid, 2015). Interestingly, the increased variation in non-CG methylome divergence in *msh1* mutants does not seem to have any significant effect on the previously described *msh1*-derived enhanced growth phenotype (Virdi et al., 2015), emphasizing the importance of *msh1*-induced CG methylation changes in this phenomenon. CG methylation changes are more stably transmitted to progeny than non-CG changes (Saze et al., 2003). A recent study also showed that non-repetitive sequences and higher CG content predispose a region for the transgenerational stability of inherited epigenetic features (Catoni et al., 2017). Moreover, stress-induced epigenetic memory is conditionally heritable through the female germline (Wibowo et al., 2016). This excludes the possibility of inheritance of stress-induced methylation changes, particularly non-CG changes to the crossed progeny of this study since *msh1* mutants were used as the pollen donor.

Results from this study indicate that *msh1* methylomes are hyper-responsive to environmental stress in a manner distinct from the wild type response, and identification of the *msh1* background as a modifier of cold-induced CHH hypomethylation provides an experimental system to further understand mechanisms that control temperature-responsive methylation changes and their inheritance behavior in crossed and selfed progeny. The experimental design of this study allowed discrimination of CG methylation changes rather than non-CG in *msh1* mutants as an influence on growth behavior of epi-lines following crossing with wild type.

## ACKNOWLEDGEMENTS

We thank Bridget Bickner for technical assistance with phenotypic measurements and PCR analysis, Dr Rebecca Roston for help in establishing freezing tolerance tests, and Dr. John Laurie for valuable conversations early in the study. We also acknowledge funding to S.M. from the Bill and Melinda Gates Foundation (0PP1088661).

## AUTHORS CONTRIBUTIONS

Conceptualization: SM and SKKR; Experiment design: SKKR and SM; Performed experiments: SKKR, with YW in biotic stress tests; Data Analysis: MRS and SKKR; Writing – Original Draft: SKKR; Writing – Review and editing: SM; All authors read and approved the final manuscript.

## SUPPLEMENTAL DATASETS

**Dataset S1:** Genes differentially expressed in *msh1* mutants that are known to respond to salt stress from Sham, et al. (2015).

**Dataset S2:** Genes differentially expressed in *msh1* mutants that are known to respond to cold and freezing temperatures from Hannah, et al. (2014).

**Dataset S3:** Hierarchical clustering of CHG-DMRs and CHH-DMRs between stressed *msh1* and wild type, with enrichment analysis for TE super families in each cluster.

**Dataset S4:** Enriched GO terms from differentially expressed genes in wild type and *msh1* mutants under different comparisons.

## SUPPLEMENT FIGURES

**Figure S1:**
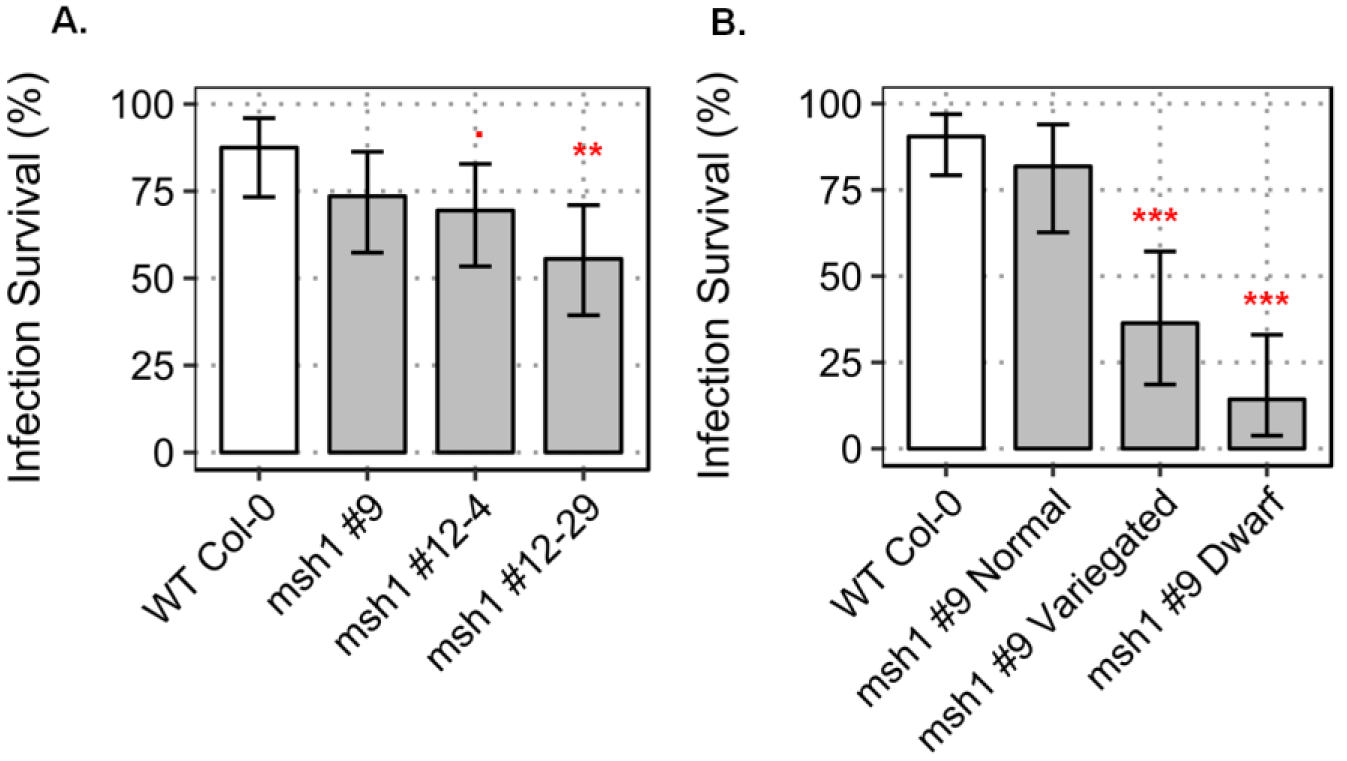
Survival rate of *A. thaliana msh1* mutants after *P. syringae* infection. A) Percent survival in *msh1* mutants and wild type after *P. syringae* pv. *tomato* DC3000 infection (n= 32 plants each). B) Survival rate of *msh1* mutants, with varying phenotype severity, and wild type after *P. syringae* pv. *tomato* DC3000 infection [for wild type n=42, *msh1* #9 n= 65 (normal phenotype =22, variegated=22, and dwarf = 21)]. Significance at '***' 0.001 '**' 0.01 '*' 0.05 '.' 0.1

**Figure S2:**
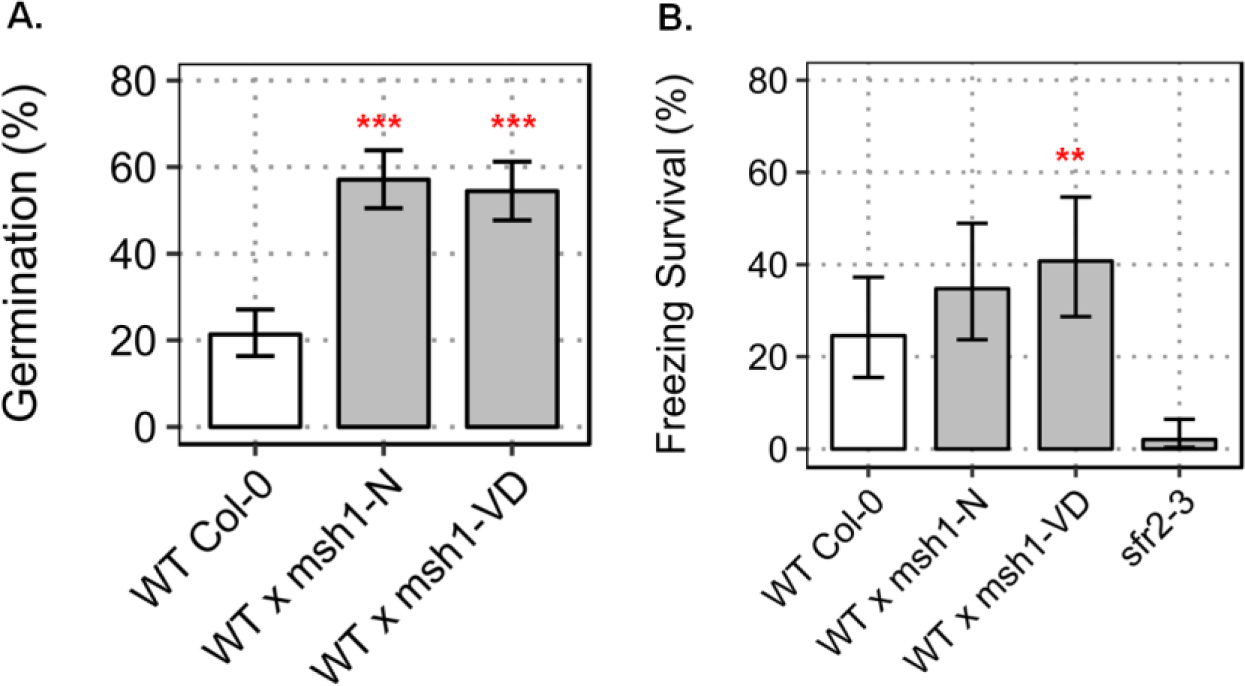
Abiotic stress tolerance in *msh1*-derived epi-lines. A) Percent germination of wild type Col-0, epi-F_3_ bulk WT x *msh1*-N and WT x *msh1*-VD at 200mM NaCl-supplemented growth media. Each bar represents three replicates (n=225), and error bars show SEM. B) Percent survival of wild type and two epi-lines after 12 hrs at -10 °C. *sjr2-3* was used as negative control. Survival was scored after one week recovery from three replicates (n=100), error bars represent SEM. Significance at '***' 0.001 '**' 0.01 '*' 0.05 '.' 0.1

**Figure S3:**
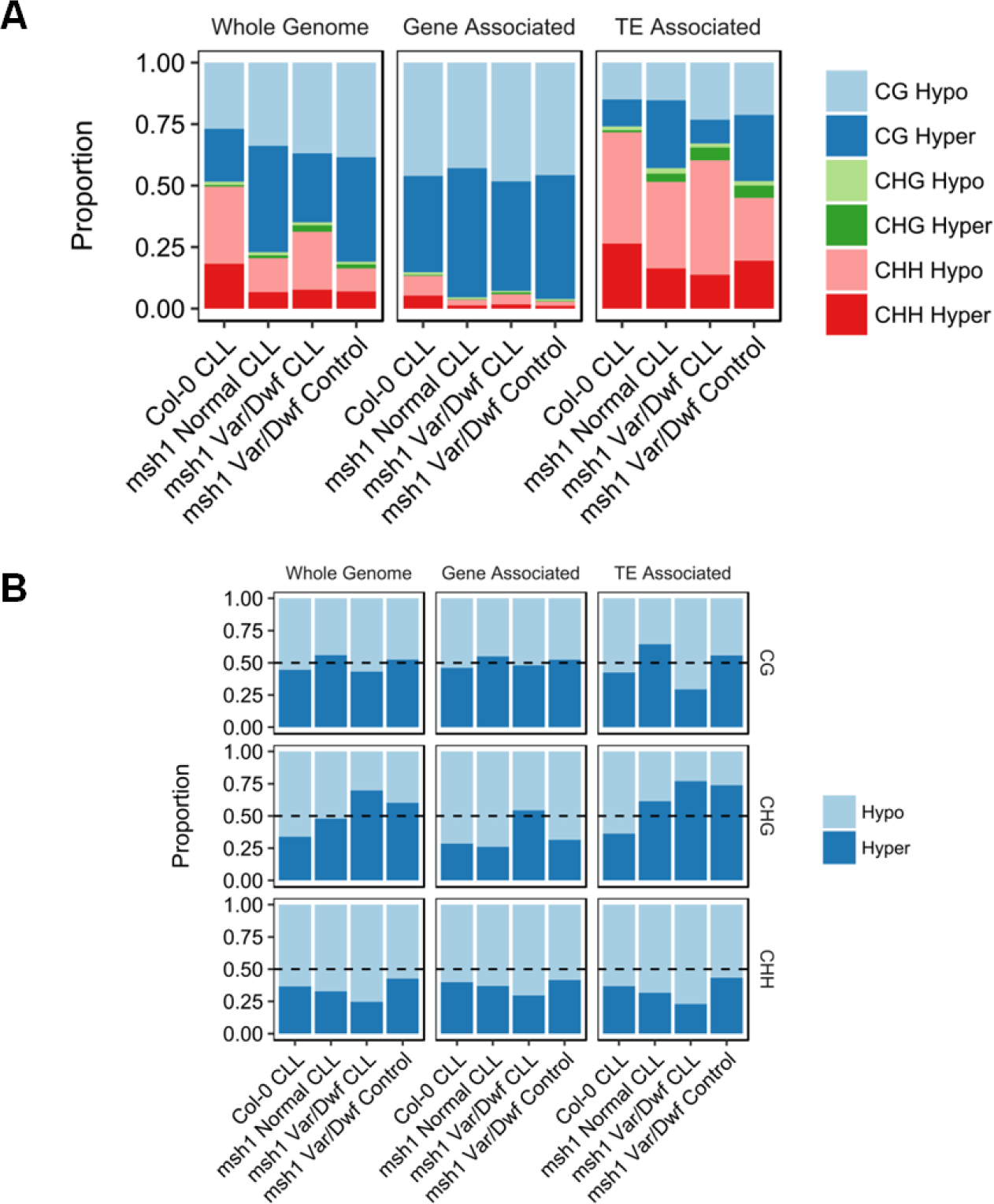
DMR distribution over genes and transposable elements in wild type and *msh1* mutants. **A)** Proportions of DMRs in each context and their genomic distribution over gene-associated and TE-associated regions of the genome. **B)** Proportion of DMRs in each cytosine context, separated based on hyper or hypomethylation. DMRs are calculated by comparing each genotype [cold-stressed wild type (Col-0 CLL) and *msh1* mutants with (CLL) or without cold stress] to control wild type Col-0.

**Figure S4:**
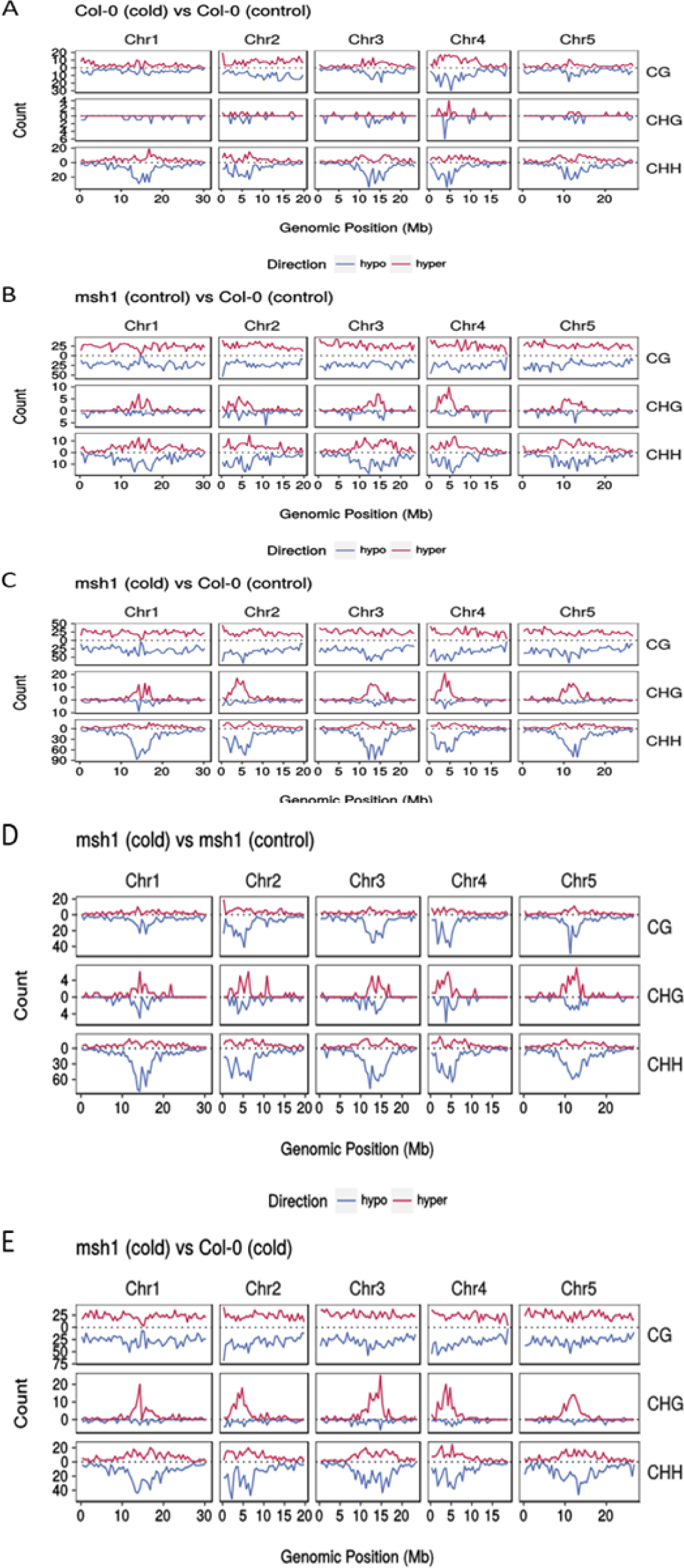
Genomic distribution of DMRs from cold treatment in *msh1* mutants. The *msh1* mutants used are variegated and dwarfed plants. **(A)** Col-0 (cold) vs Col-0 (control). **(B)** *msh1* (control) vs Col-0 (control). **(C)** *msh1* (cold) vs Col-0 (control). **(D)** *msh1* (cold) vs *msh1* (control). **(E)** *msh1* (cold) vs Col-0 (control). Note the y-axes are independently scaled by comparison and context.

**Figure S5:**
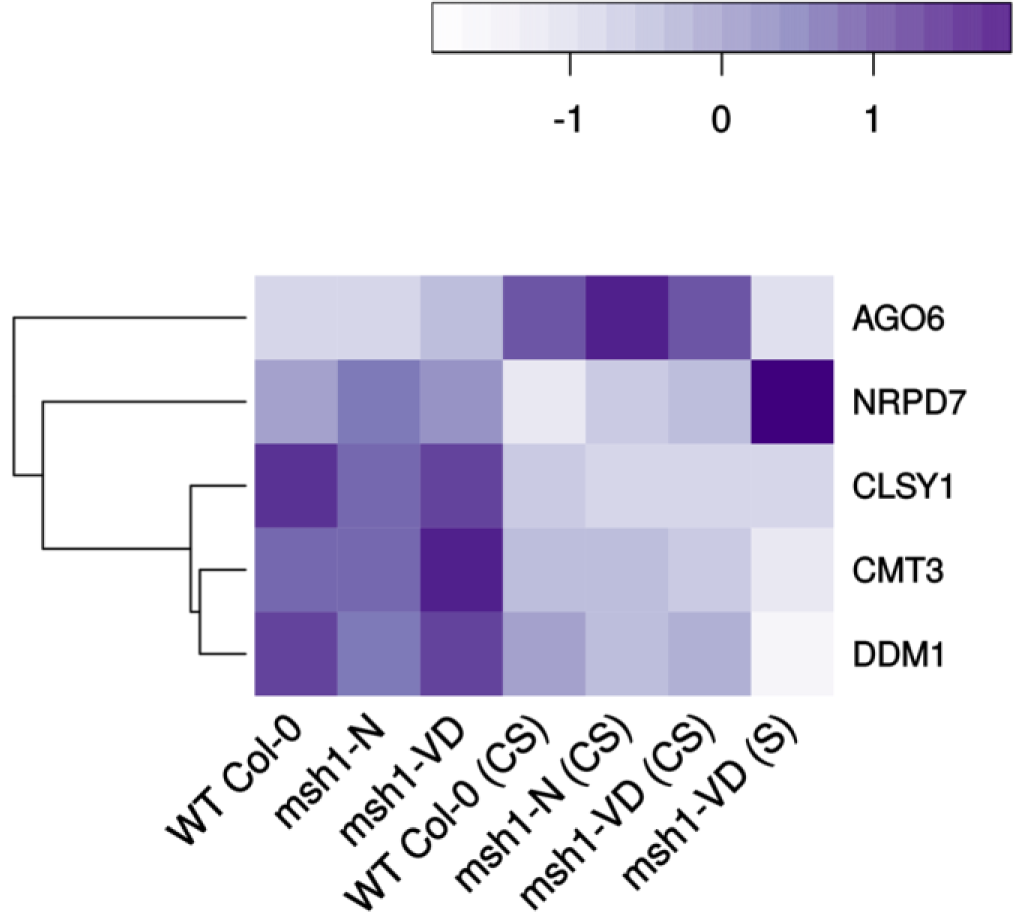
Heatmap showing differential expression of methylation machinery genes in *msh1* mutants under chronic cold stress. Genes differentially expressed (normalized fold change) in at least one comparison between wild type and *msh1* mutants were considered for the heatmap, from a list of genes involved in methylation machinery (Matzke, et al. 2009).

**Figure S6:**
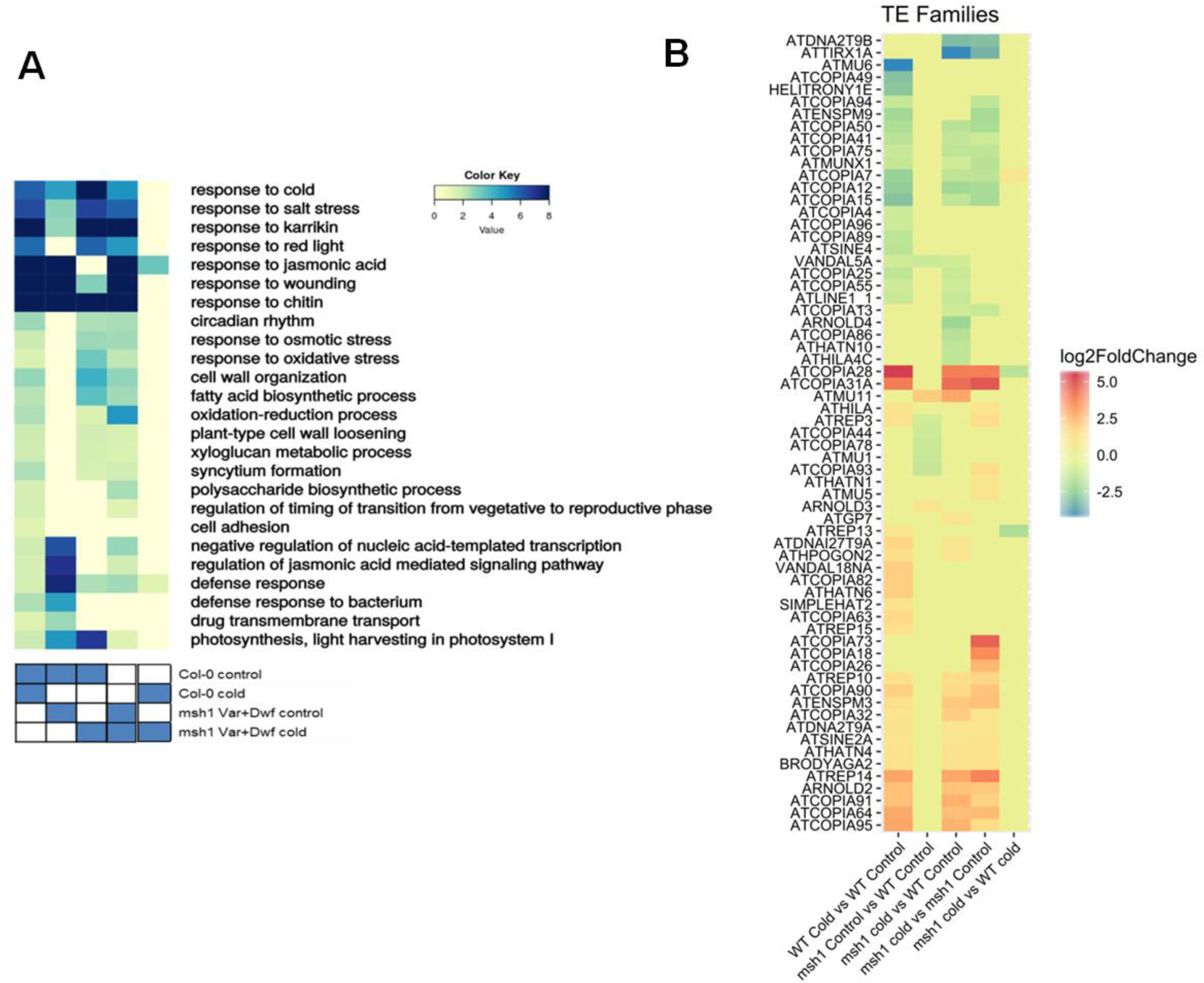
Heat maps showing differential expression of genes and transposable elements. **A)** Heat map showing enriched GO terms for differentially expressed genes in comparisons between wild type and *msh1* mutants with or without stress. **B)** Heat map showing differential expression of transposable elements at the family level.

**Figure S7.**
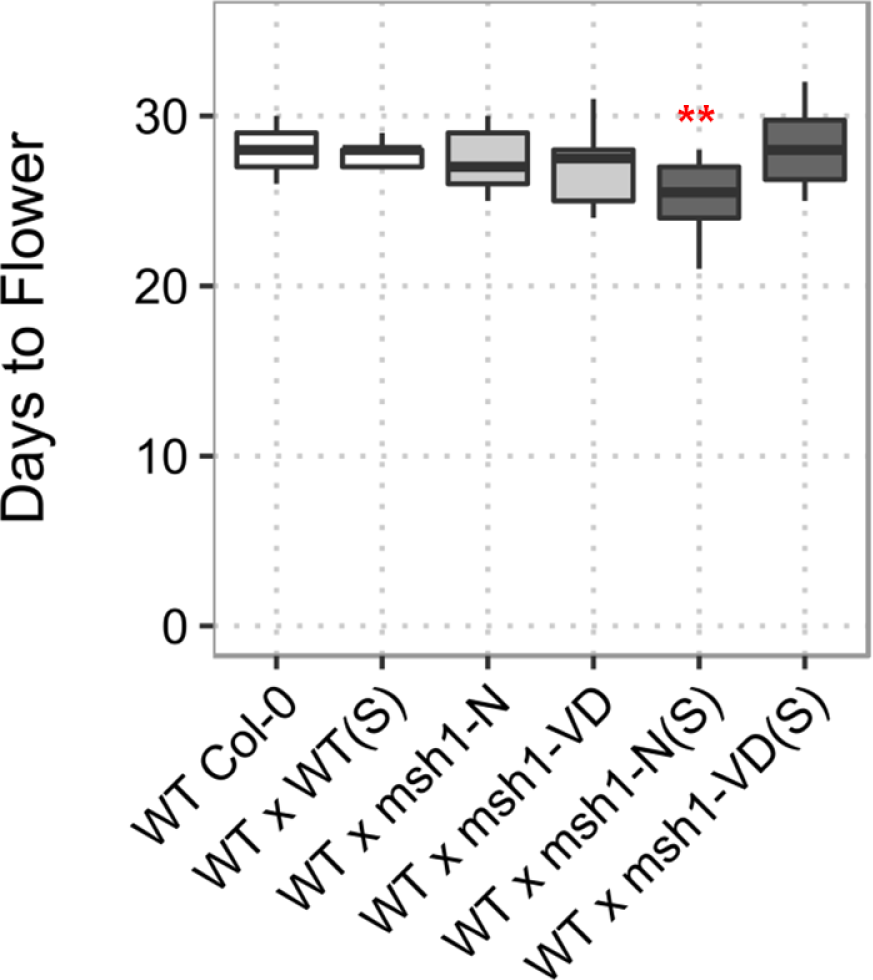
Variation in days to flowering in epi-F_2s_ derived from cold-stressed (S) and unstressed *msh1* mutants. Days to flowering was measured as number of days from germination to the first open flower. Eighteen plants were sampled for each F2 population and wild type. Plants were grown in 12/12 hr light/dark cycle at 22 °C. Significance at '***' 0.001 '**' 0.01 '*' 0.05 '.' 0.1.

## REFERENCES

Akalin, A., Kormaksson, M., Li, S., Garrett-Bakelman, F.E., Figueroa, M.E., Melnick, A. and Mason, C.E. (2012) methylKit: a comprehensive R package for the analysis of genome-wide DNA methylation profiles. Genome biology 13, R87.

Barnes, A.C., Benning, C. and Roston, R. (2016) Chloroplast membrane remodeling during freezing stress is accompanied by cytoplasmic acidification activating SENSITIVE TO FREEZING 2. Plant physiology, pp. 00286.02016.

Benjamini, Y. and Hochberg, Y. (1995) Controlling the false discovery rate: a practical and powerful approach to multiple testing. Journal of the royal statistical society. Series B (Methodological), 289–300.

Bilichak, A. and Kovalchuk, I. (2016) Transgenerational response to stress in plants and its application for breeding. J Exp Bot 67, 2081–2092.

Cao, S., Ye, M. and Jiang, S. (2005) Involvement of GIGANTEA gene in the regulation of the cold stress response in Arabidopsis. Plant cell reports 24, 683–690.

Catoni, M., Griffiths, J., Becker, C., Zabet, N.R., Bayon, C., Dapp, M., Lieberman-Lazarovich, M., Weigel, D. and Paszkowski, J. (2017) DNA sequence properties that predict susceptibility to epiallelic switching. The EMBO Journal.

Chow, H.T. and Ng, D.W. (2017) Regulation of miR163 and its targets in defense against Pseudomonas syringae in Arabidopsis thaliana. Scientific Reports 7, 46433.

de la Rosa Santamaria, R., Shao, M.R., Wang, G., Nino-Liu, D.O., Kundariya, H., Wamboldt, Y., Dweikat, I. and Mackenzie, S.A. (2014) MSH1-induced non-genetic variation provides a source of phenotypic diversity in Sorghum bicolor. PLoS One 9, e108407.

Dobin, A., Davis, C.A., Schlesinger, F., Drenkow, J., Zaleski, C., Jha, S., Batut, P., Chaisson, M. and Gingeras, T.R. (2013) STAR: ultrafast universal RNA-seq aligner. Bioinformatics 29.

Dong, M.A., Farré, E.M. and Thomashow, M.F. (2011) Circadian clock-associated 1 and late elongated hypocotyl regulate expression of the C-repeat binding factor (CBF) pathway in Arabidopsis. Proceedings of the National Academy of Sciences 108, 7241–7246.

Dowen, R.H., Pelizzola, M., Schmitz, R.J., Lister, R., Dowen, J.M., Nery, J.R., Dixon, J.E. and Ecker, J.R. (2012) Widespread dynamic DNA methylation in response to biotic stress. Proc Natl Acad Sci U S A 109, E2183–2191.

Dubin, M.J., Zhang, P., Meng, D., Remigereau, M.-S., Osborne, E.J., Casale, F.P., Drewe, P., Kahles, A., Jean, G. and Vilhjálmsson, B. (2015) DNA methylation in Arabidopsis has a genetic basis and shows evidence of local adaptation. Elife 4, e05255.

Franks, S.J. and Hoffmann, A.A. (2012) Genetics of climate change adaptation. Annu Rev Genet 46, 185–208.

Hannah, M.A., Wiese, D., Freund, S., Fiehn, O., Heyer, A.G. and Hincha, D.K. (2006) Natural genetic variation of freezing tolerance in Arabidopsis. Plant physiology 142, 98–112.

Hsu, P.Y. and Harmer, S.L. (2012) Circadian phase has profound effects on differential expression analysis. PLoS One 7, e49853.

Huang, D.W., Sherman, B.T. and Lempicki, R.A. (2009) Systematic and integrative analysis of large gene lists using DAVID bioinformatics resources. Nature protocols 4, 44–57.

Jin, Y., Tam, O.H., Paniagua, E. and Hammell, M. (2015) TEtranscripts: a package for including transposable elements in differential expression analysis of RNA-seq datasets. Bioinformatics 31.

Kim, D., Pertea, G., Trapnell, C., Pimentel, H., Kelley, R. and Salzberg, S.L. (2013) TopHat2: accurate alignment of transcriptomes in the presence of insertions, deletions and gene fusions. Genome Biol 14.

King, E.O., Ward, M.K. and Raney, D.E. (1954) Two simple media for the demonstration of pyocyanin and fluorescin. Journal of laboratory and clinical medicine 44, 301–307.

Kinoshita, T. and Seki, M. (2014) Epigenetic memory for stress response and adaptation in plants. Plant Cell Physiol 55.

Krueger, F. and Andrews, S.R. (2011) Bismark: a flexible aligner and methylation caller for Bisulfite-Seq applications. bioinformatics 27, 1571–1572.

Lewsey, M.G., Hardcastle, T.J., Melnyk, C.W., Molnar, A., Valli, A., Urich, M.A., Nery, J.R., Baulcombe, D.C. and Ecker, J.R. (2016) Mobile small RNAs regulate genome-wide DNA methylation. Proceedings of the National Academy of Sciences 113, E801–E810.

Li, J. and Chory, J. (1998) Preparation of DNA from Arabidopsis. Arabidopsis Protocols, 55–60.

Lockton, S. and Gaut, B.S. (2009) The contribution of transposable elements to expressed coding sequence in Arabidopsis thaliana. Journal of Molecular Evolution 68, 80–89.

Love, M.I., Huber, W. and Anders, S. (2014) Moderated estimation of fold change and dispersion for RNA-seq data with DESeq2. Genome Biol 15.

Maibam, P., Nawkar, G.M., Park, J.H., Sahi, V.P., Lee, S.Y. and Kang, C.H. (2013) The influence of light quality, circadian rhythm, and photoperiod on the CBF-mediated freezing tolerance. International journal of molecular sciences 14, 11527–11543.

Moellering, E.R., Muthan, B. and Benning, C. (2010) Freezing tolerance in plants requires lipid remodeling at the outer chloroplast membrane. Science 330, 226–228.

Novillo, F., Medina, J. and Salinas, J. (2007) Arabidopsis CBF1 and CBF3 have a different function than CBF2 in cold acclimation and define different gene classes in the CBF regulon. Proceedings of the National Academy of Sciences 104, 21002–21007.

Probst, A.V. and Mittelsten Scheid, O. (2015) Stress-induced structural changes in plant chromatin. Curr Opin Plant Biol 27.

Raju, S.K.K., Shao, M.-R., Sanchez, R., Xu, Y.-Z., Sandhu, A., Graef, G. and Mackenzie, S. (2017) An epigenetic breeding system in soybean for increased yield and stability. bioRxiv, 232819.

Sanchez, R., Yang, X., Kundariya, H., Barreras, J.R., Wamboldt, Y. and Mackenzie, S. (2018) Enhancing resolution of natural methylome reprogramming behavior in plants. bioRxiv, 252106.

Saze, H., Scheid, O.M. and Paszkowski, J. (2003) Maintenance of CpG methylation is essential for epigenetic inheritance during plant gametogenesis. Nature genetics 34, 65–69.

Sham, A., Moustafa, K., Al-Ameri, S., Al-Azzawi, A., Iratni, R. and AbuQamar, S. (2015) Identification of Arabidopsis candidate genes in response to biotic and abiotic stresses using comparative microarrays. PLoS One 10.

Shao, M.-R., Kumar Kenchanmane Raju, S., Laurie, J.D., Sanchez, R. and Mackenzie, S.A. (2017) Stress-responsive pathways and small RNA changes distinguish variable developmental phenotypes caused by MSH1 loss. BMC Plant Biology 17, 47.

Shedge, V., Davila, J., Arrieta-Montiel, M.P., Mohammed, S. and Mackenzie, S.A. (2010) Extensive rearrangement of the Arabidopsis mitochondrial genome elicits cellular conditions for thermotolerance. Plant physiology 152, 1960–1970.

Slotkin, R.K. and Martienssen, R. (2007) Transposable elements and the epigenetic regulation of the genome. Nat Rev Genet 8.

Stroud, H., Do, T., Du, J., Zhong, X., Feng, S., Johnson, L., Patel, D.J. and Jacobsen, S.E. (2014) Non-CG methylation patterns shape the epigenetic landscape in Arabidopsis. Nature structural & molecular biology 21, 64–72.

Stroud, H., Greenberg, M.V., Feng, S., Bernatavichute, Y.V. and Jacobsen, S.E. (2013) Comprehensive analysis of silencing mutants reveals complex regulation of the Arabidopsis methylome. Cell 152, 352–364.

Uller, T., English, S. and Pen, I. (2015) When is incomplete epigenetic resetting in germ cells favoured by natural selection? In: Proc. R. Soc. B p. 20150682. The Royal Society.

Virdi, K.S., Laurie, J.D., Xu, Y.Z., Yu, J., Shao, M.R., Sanchez, R., Kundariya, H., Wang, D., Riethoven, J.J., Wamboldt, Y., Arrieta-Montiel, M.P., Shedge, V. and Mackenzie, S.A. (2015) Arabidopsis MSH1 mutation alters the epigenome and produces heritable changes in plant growth. Nat Commun 6, 6386.

Virdi, K.S., Wamboldt, Y., Kundariya, H., Laurie, J.D., Keren, I., Kumar, K.S., Block, A., Basset, G., Luebker, S. and Elowsky, C. (2016) MSH1 is a plant organellar DNA binding and thylakoid protein under precise spatial regulation to alter development. Molecular Plant 9, 245–260.

Ward Jr, J.H. (1963) Hierarchical grouping to optimize an objective function. Journal of the American statistical association 58, 236–244.

Wessler, S.R. (1996) Plant retrotransposons: turned on by stress. Current Biology 6, 959–961.

Wibowo, A., Becker, C., Marconi, G., Durr, J., Price, J., Hagmann, J., Papareddy, R., Putra, H., Kageyama, J. and Becker, J. (2016) Hyperosmotic stress memory in Arabidopsis is mediated by distinct epigenetically labile sites in the genome and is restricted in the male germline by DNA glycosylase activity. Elife 5, e13546.

Xu, Y.Z., Arrieta-Montiel, M.P., Virdi, K.S., de Paula, W.B., Widhalm, J.R., Basset, G.J., Davila, J.I., Elthon, T.E., Elowsky, C.G., Sato, S.J., Clemente, T.E. and Mackenzie, S.A. (2011) MutS HOMOLOG1 is a nucleoid protein that alters mitochondrial and plastid properties and plant response to high light. Plant Cell 23, 3428–3441.

Xu, Y.Z., Santamaria Rde, L., Virdi, K.S., Arrieta-Montiel, M.P., Razvi, F., Li, S., Ren, G., Yu, B., Alexander, D., Guo, L., Feng, X., Dweikat, I.M., Clemente, T.E. and Mackenzie, S.A. (2012) The chloroplast triggers developmental reprogramming when mutS HOMOLOG1 is suppressed in plants. Plant Physiol 159, 710–720.

Yang, X., Kundariya, H., Xu, Y.Z., Sandhu, A., Yu, J., Hutton, S.F., Zhang, M. and Mackenzie, S.A. (2015) MutS HOMOLOG1-derived epigenetic breeding potential in tomato. Plant Physiol 168, 222–232.

Yong-Villalobos, L., González-Morales, S.I., Wrobel, K., Gutiérrez-Alanis, D., Cervantes-Peréz, S.A., Hayano-Kanashiro, C., Oropeza-Aburto, A., Cruz-Ramírez, A., Martínez, O. and Herrera-Estrella, L. (2015) Methylome analysis reveals an important role for epigenetic changes in the regulation of the Arabidopsis response to phosphate starvation. Proc Natl Acad Sci U S A 112.

Zilberman, D. (2017) An evolutionary case for functional gene body methylation in plants and animals. Genome biology 18, 87.

